# A simple and efficient metric quantifying druggable property of chemical small molecules

**DOI:** 10.1101/2020.07.13.199752

**Authors:** Chuanbo Huang, Jichun Yang, Qinghua Cui

## Abstract

One big class of drugs are chemical small molecules (CSMs), but the majority of CSMs are in very low druggable potential. Therefore, it is quite important to predict drug-related properties (druggable properties) for candidate CSMs. Currently, although a number of druggable properties (e.g. logP and pKa) can be calculated by in silico methods, the identification of druggable CSMs is still at high risk and new quantitative metrics for the druggable potential of CSMs are increasingly needed. Here, we presented normalized bond energy (NBE), a new metric for the above purpose. By applying NBE to the DrugBank CSMs whose properties are largely known, we revealed that NBE is able to describe a number of critical druggable properties including logP, pKa, membrane permeability, blood-brain barrier penetration, and human intestinal absorption. Moreover, given that the human endogenous metabolites could be served as an important resource for drug discovery, we applied NBE to the metabolites in the Human Metabolome Database. As a result, NBE shows a significant difference in metabolites from various body fluids and is correlated with some important properties including melting point and water solubility.

## Introduction

Research and development of pharmaceuticals is a resource-consuming and long process with a variety of challenging risks[1]. Chemical small molecules (CSMs) represent a big class of drugs which mainly function by binding with disease related target molecules [2]. Given the huge space of target molecules and CSMs, evaluating the druggable potential of both targets [3–6] and CSMs [7–11] is thus one of the hearts of drug discovery. For CSMs, it is known that a number of drug related properties (druggable properties) affect their druggable potential, for example human intestinal absorption, blood-brain barrier penetration[12], and some pharmacokinetic properties[13]. Therefore, it is important to accurately predicted druggable properties for early phase candidate CSM and large-scale druggable CSM screening. For the above purpose, a number of in-silico methods or metrics have already been proposed. For example, properties of logP, logD, logS, logW, and pKa can be calculated using the free online tool ALOGPS[14]. ChemAxon, a tool providing solutions and services for chemistry and biology[15], can be used to predict druggable properties such as water solubility, polar surface area (PSA), H bond acceptor count, H bond donor count, and pKa etc. In addition, given that CSM transport is a key factor for drug potential, several metrics have been presented to evaluate CSM transport properties, including octanol/water partition coefficient, molecular size and shape, hydrogen-bonding capabilities and topological polar surface area[16–19]. These in-silico methods or metrics did provide helps for quickly quantifying CSM properties in order to evaluate their druggable potential. However, due to the huge complexity of both biology and chemistry, these methods or metrics are still far from solving all problems in drug research and development, which is still in high risks. For example, when chemical structures are diverse, the molecular transport related physicochemical metrics introduced above descriptors will be not reliable to predict molecular transport properties[20]. Thus it is necessary to present new in-silico methods or metrics to quantify druggable properties of CSMs.

We previously revealed that the free energy of RNA secondary structure has a significant contribution to the importance score of both protein-coding RNA molecules (mRNAs) and noncoding RNAs (lncRNAs and miRNAs)[21,22]. Based on the above observations, it is hypothesized that the energy status of CSMs could represent some property of these molecules. To confirm this hypothesis, here we presented a new metric, normalized bond energy (NBE). Moreover, we revealed that NBE score can significantly represent some critical druggable properties, such as logP, pKa, permeability, blood-brain barrier penetration, and human intestinal absorption. Moreover, given that the human endogenous metabolites could be served as a resource for drug discovery[23], we calculated the NBE scores for CSMs in the human metabolome and performed a comprehensive bioinformatic analysis for the relations between NBE score and other properties of these endogenous metabolic small molecules.

## Materials and Methods

### Datasets of CSMs

We obtained the structure data in SDF format of CSMs from the Drugbank database[2](Version 5.0) which includes approved small molecule drugs and experimental drugs. We obtained the structure data in SDF format of small molecule metabolites from the Human Metabolome Database (HMDB)[24]. Experiment-derived property (e.g. melting point, logP, pKa, etc) data of CSMs were also curated from DrugBank and HMDB. For the property of water solubility, CSMs with terms like “Insoluble”, “almost insoluble”, “Low”, “Mostly insoluble”, “Non-soluble”, “not soluble”, and “Poorly soluble” etc were assigned as the insoluble group; whereas CSMs with terms like “Soluble”, “Easily soluble”, “Complete”, “Freely soluble”, “Highly soluble”, and “Very soluble” etc were assigned as the soluble group. In addition, we obtained the Caco-2 monolayers permeability data of 690 CSMs from the study by Waterbeemd et al.[18,25,26], the blood-brain barrier penetration data of 1638 CSMs from the study by Kelder et al.[27,28], and the human intestinal absorption data of 598 CSMs from the study by Shen et al[28,29]. The structure files of these CSMs in SMILES format were obtained as well.

### Calculation of normalized bond energy

For one CSM, we first extracted its bonds from its molecular structure using the RDKit library[30] (version 2017.09.1, 2017), and then determine the energy (KJ/mol)of each bond by matching it with the bond energy table (Supplement Table 1)through the bond type and the atom type. The bond type here includes single bond, double bond and triple bond, which is the number of shared electron pairs between the two corresponding atoms. Bond energy is defined to physically quantify the strength of a chemical bond and can be measured by the amount of energy required when the bond breaks, and it is an average value of bond dissociation energy of a mole of molecule in the gas phase, typically at a temperature of 298 Kelvin. Given that bigger molecules normally have more bonds and thus would have bigger bond energy, we next defined normalized bond energy (NBE), which normalizes the original bond energy using molecular weight calculated using RDKit. The procedure of the algorithm was shown in Figure 1.

**Figure 1.**
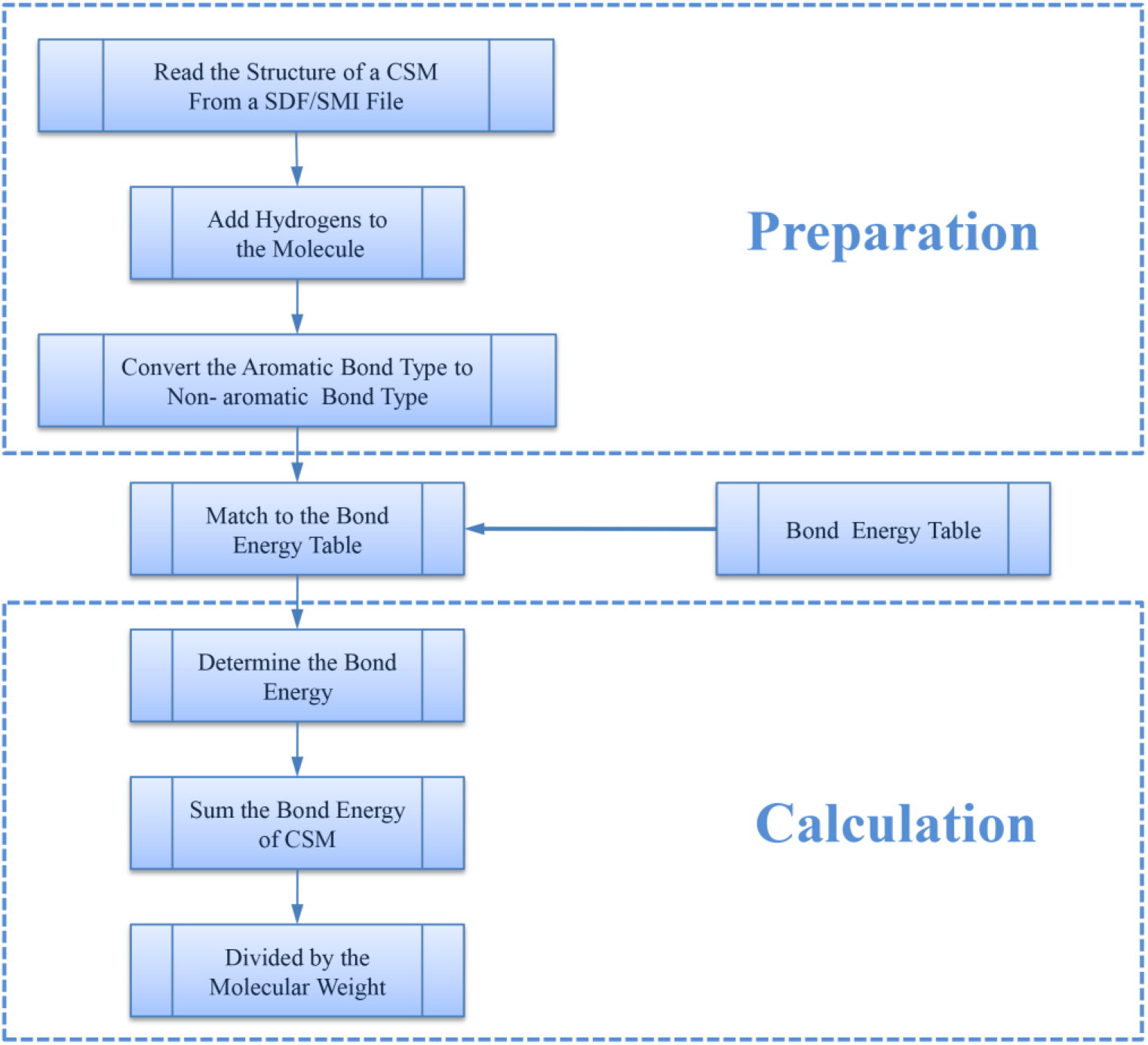
The flowchart of NBE algorithms.

Then, NBE can be calculated using the following equation.

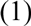

Where is the bond energy of bond, is the number of bonds, and is the molecular weight of the CSM.

### Statistical computation

We implemented the algorithm of NBE (http://www.cuilab.cn/nbe/nbe.zip) using python. Spearman’s correlation analysis, t test and Wilcoxon test were performed using R.

## Results

### Global distribution of NBE scores

The whole framework of this study is shown as Figure 2. As a result, we calculated NBE scores for 10426 CSMs (2444are approved drugs and 7982 are experimental drugs) from DrugBank and those for 113878 human metabolic CSMs from HMDB. The distributions of the DrugBank NBE scores are shown as Figure 3A. The approved drugs have greater NBE scores than the unapproved drugs (p-value=1.17e-13, Wilcoxon test; Figure 3B).

**Figure 2.**
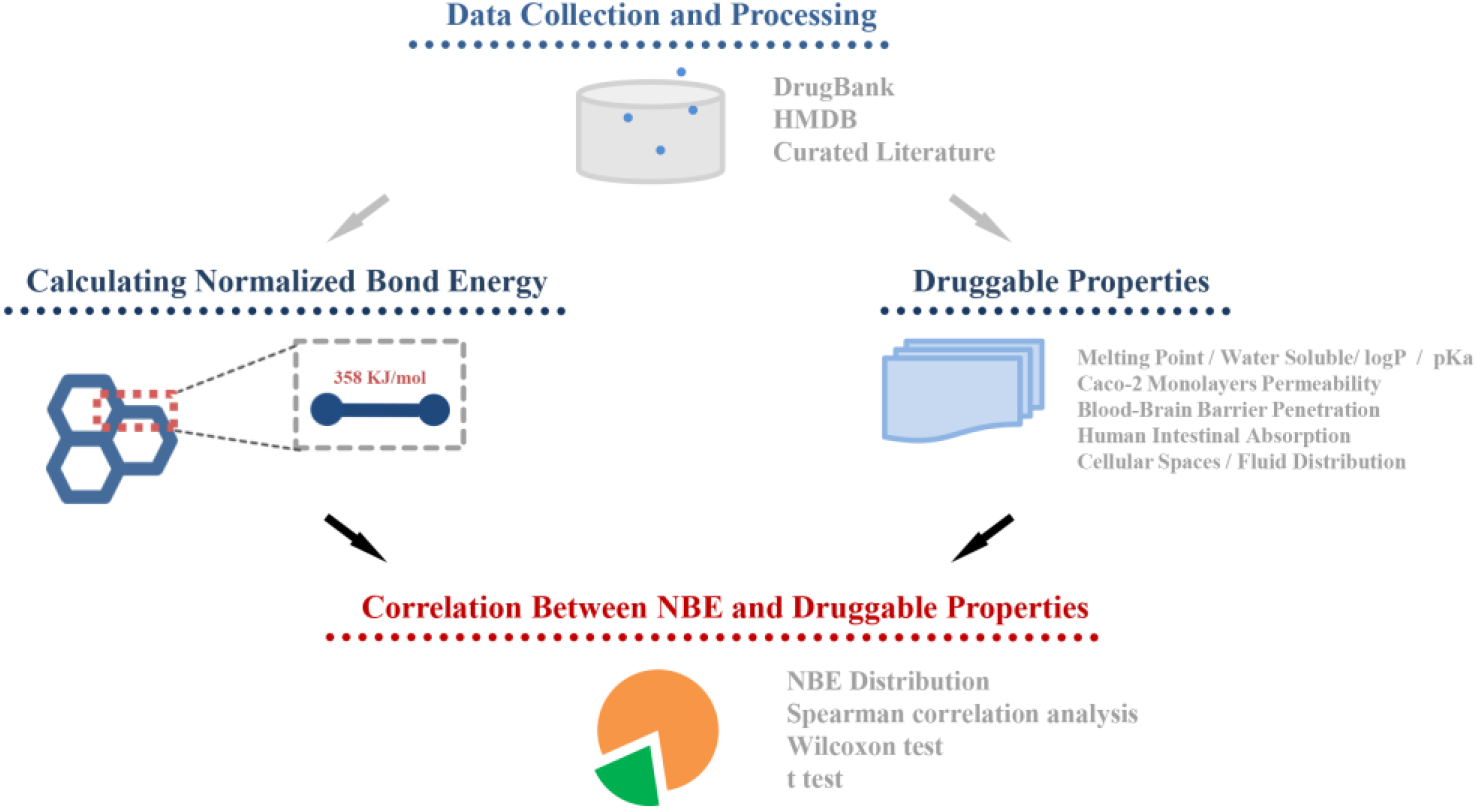
The framework of the whole study.

**Figure 3.**
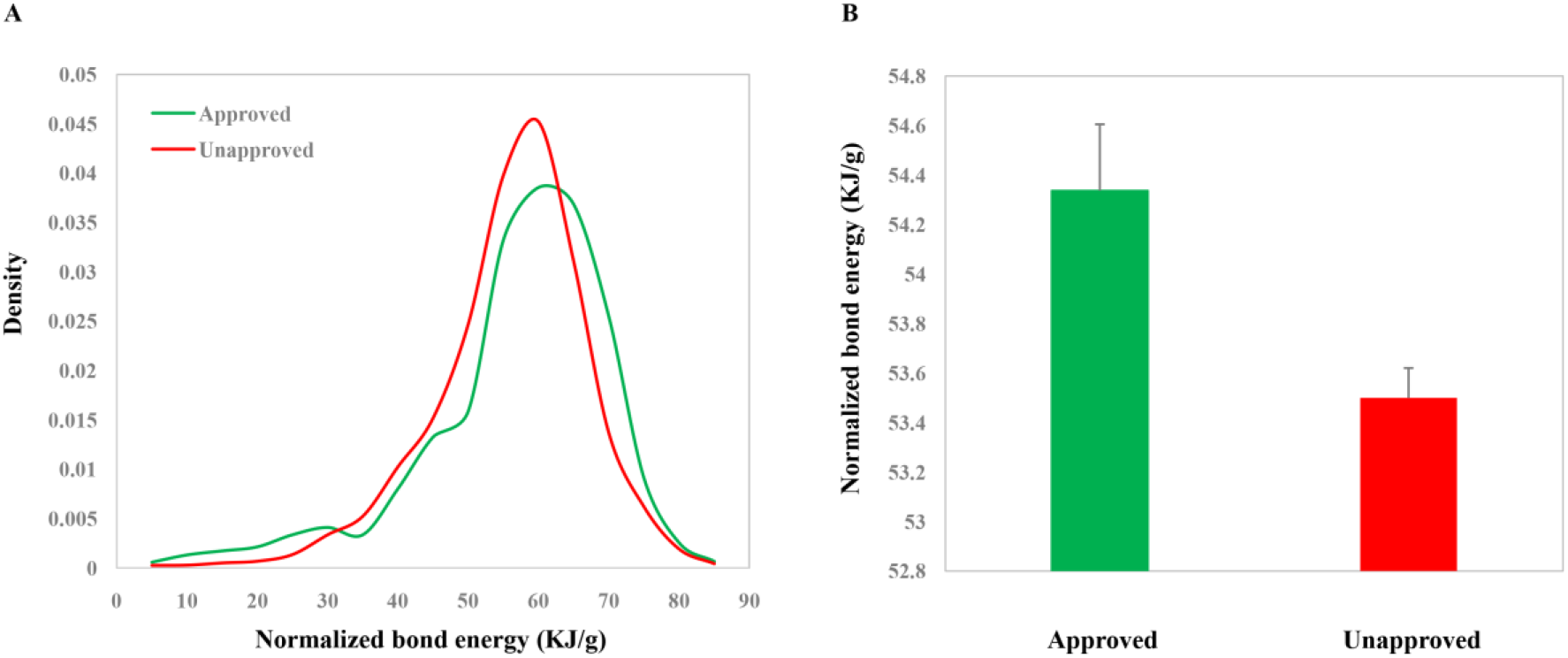
Distributions (A) and values (B) of the NBE scores of approved chemical small molecules (CSMs) and unapproved ones in DrugBank.

### Correlations of NBE scores with experimentally identified properties of CSMs in DrugBank

As a result, we found that NBE score is significantly associated with a number of experimentally identified properties of CSMs in DrugBank. Firstly, we asked whether there is difference in NBE scores between the soluble CSMs and the insoluble ones. The result showed that the soluble CSMs have smaller NBE scores than the insoluble ones (mean: 53.3 vs. 56.7, p-value=0.009, t test; Figure 4A). Moreover, we investigated the relations of NBE score with melting point, logP, and pKa. We found that NBE score shows significantly negative correlation with melting point (Rho=−0.19, p-value=1.89e-13, Figure 4B) but shows positive correlation with logP (Rho=0.36, p-value=1.25e-47, Figure 4C) and pKa (Rho=0.38, p-value=9.08e-19, Figure 4D).

**Figure 4.**
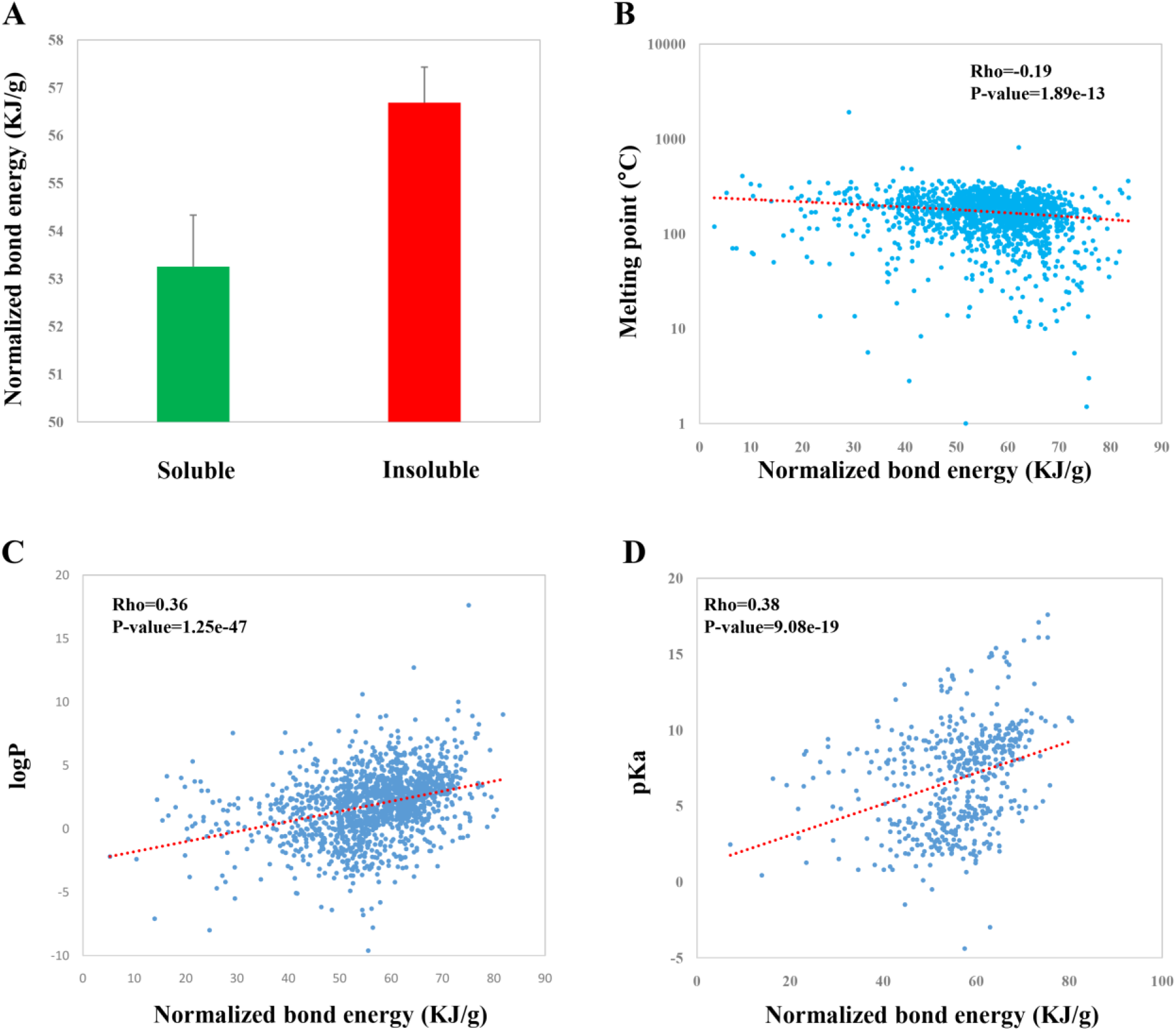
Correlations of the NBE scores of chemical small molecules (CSMs) in DrugBank with druggable properties including water solubility (A), melting point (B), logP (C), and pKa (D).

### NBE is correlated with permeability, blood-brain barrier penetration, and human intestinal absorption

Permeability, blood-brain barrier penetration, and human intestinal absorption are another three critical properties which greatly affect the transport properties of a CSM. We observed significant correlations between NBE and Caco-2 monolayers permeability (Rho=0.80,p-value=0.01, Figure 5A;Rho=0.71, p-value=0.003, Figure 5B; Rho=0.22, p-value=9.186e-09, Figure 5C), between NBE and human intestinal absorption (Rho=0.44, p-value=0.05, Figure 5D, Rho=0.22, p-value=3.567e-03, Figure 5E), and between NBE and blood-brain barrier penetration (Rho=0.55,p-value=9.39e-05, Figure 5F). Moreover, using the FA% values threshold of 30%, we divide CSMs of human intestinal absorption dataset into the absorbable(HIA+)terms and the unabsorbable(HIA−)terms. We observed that HIA+ CSMs have bigger NBE scores than the HIA-CSMs (mean: 57.0 vs. 52.4, p-value=0.0064, t test; Figure 5G). And we also observed that brain-blood barrier penetratable (BBB+) CSMs have bigger NBE scores than the unpenetratable (BBB−) CSMs (mean: 58.7 vs. 54.2, p-value=1.08e-15, t test; Figure 5H).

**Figure 5.**
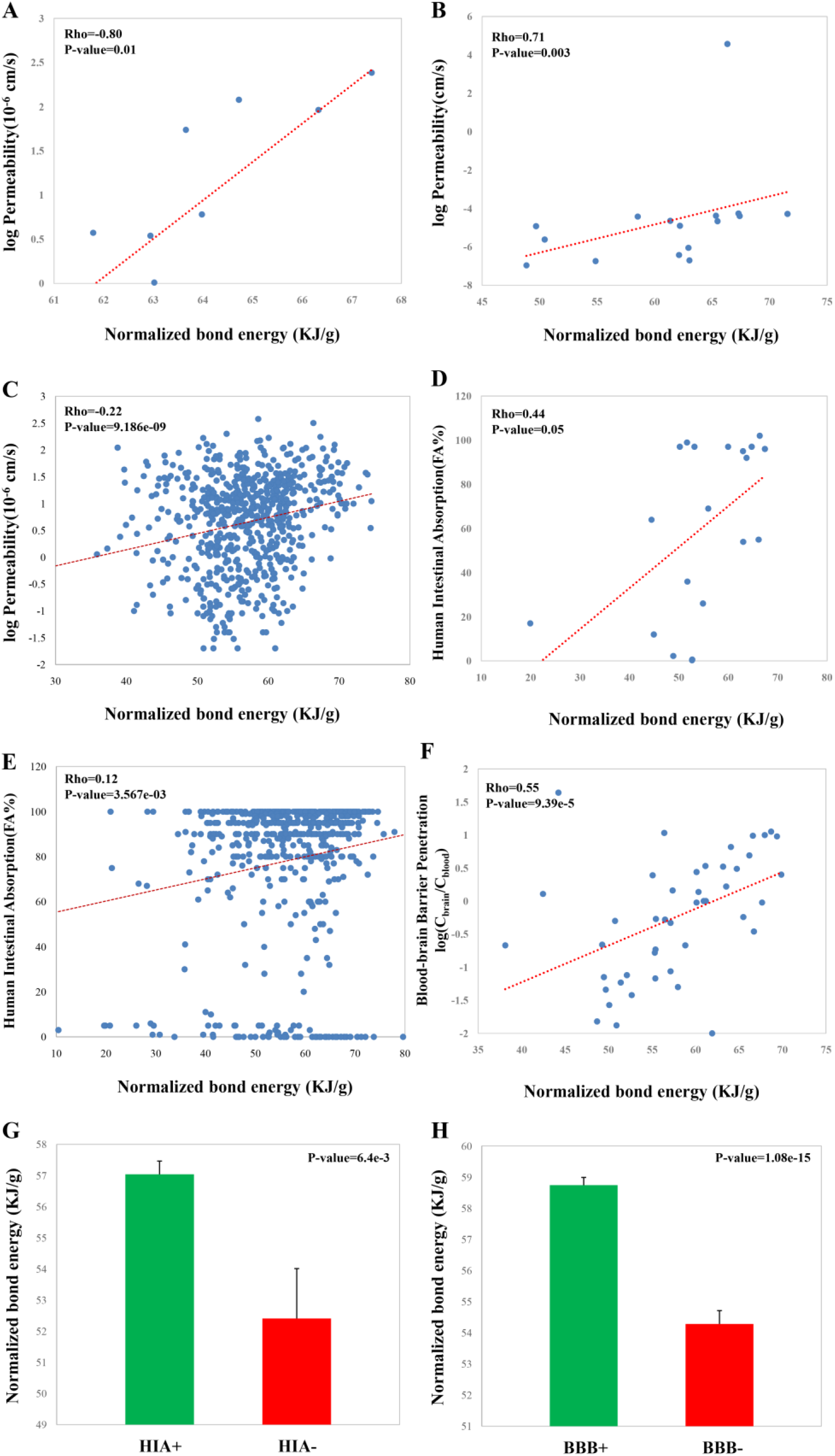
**Correlations of the NBE scores of chemical small molecules (CSMs) with some druggable properties,** including Caco-2 monolayers permeability from reference[18] (A), reference[25] (B) and reference[26] (C), human intestinal absorption from reference[29] (D) and reference[28] (E), blood-brain barrier penetration from reference[27] (F). Values of the NBE scores of human intestinal absorbable (HIA+) CSMs and unabsorbable (HIA-) ones from reference[28](G).Values of the NBE scores of brain-blood barrier penetratable (BBB+) CSMs and unpenetratable (BBB−) ones from reference[27] (H).

### NBE is correlated with properties of human metabolites

Natural products represent one important class of CSMs with high druggable potential. As one type of natural products, the human endogenous metabolites could be served as a resource for drug discovery. It is thus important to investigate whether NBE can describe some properties of human metabolites. We previously revealed biased subcellular distributions for miRNA target genes and sex-biased genes. We found that miRNAs prefer to target genes located within inner cellular space compared to genes located in outer cellular space[31]. Female biased genes are enriched in outer cellular space; whereas male biased genes are enriched in inner cellular space [32]. It thus will be interesting to investigate whether there exist difference in NBE for metabolites in different cellular spaces. The result showed that metabolites in outer cellular space (metabolites in extracellular space and/or membrane) have greater NBE scores than those in inner cellular space (metabolites in cytoplasm and/or nucleus) (p-value=0, Wilcoxon test; Figure 6A). Moreover, metabolites derived from different body fluids show significant different NBE scores (p-value=0.0, spearman test, Figure 6B). Metabolites from feces and saliva show the highest NBE scores (Figure 6B). In addition, NBE scores of metabolites are correlated with melting point (Rho=−0.29, p-value=7.11e-138; Figure 6C) and water solubility (Rho=−0.29, p-value=1.11e-51; Figure 6D), which is consistent with the results on CSMs in DrugBank.

**Figure. 6.**
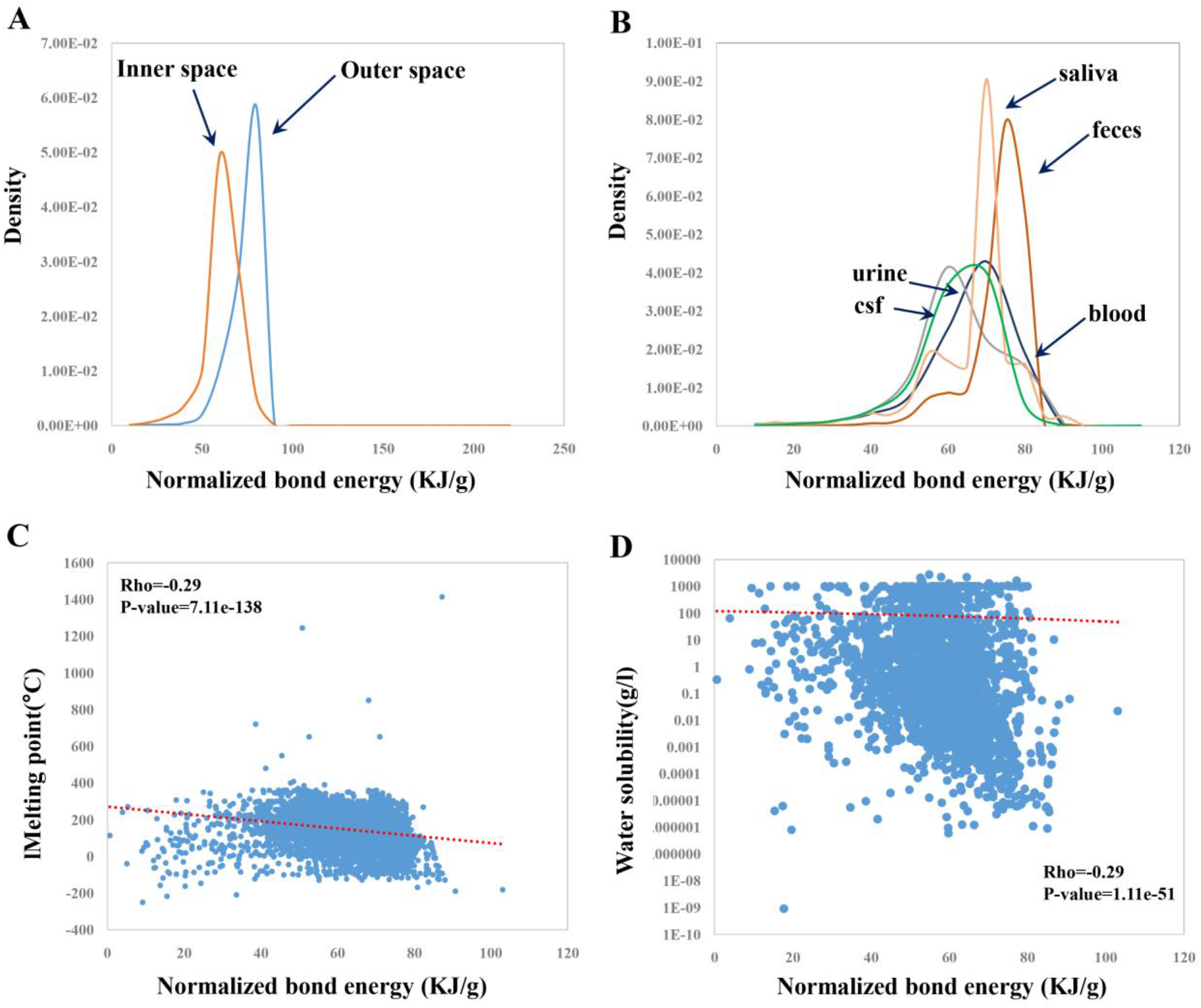
Distributions of the NBE scores of metabolites in different cellular locations (A) and of metabolites from different body fluids (B), and correlation of the NBE scores with melting point (C) and with water solubility (D) in HMDB.

## Discussion

In this study, we presented a new in-silico metric, normalized bond energy (NBE), for quantifying some property of chemical small molecules (CSMs). The results showed that NBE is able to describe some critical druggable properties of CSMs, for example blood-brain barrier penetration and human intestinal absorption.

The blood-brain barrier separates the brain from the systemic blood circulation and maintains the homeostasis of the central nervous system. Thus the blood-brain distribution of a CSM is a key characteristic for determing whether it is potentially druggable for the central nervous system. Human intestinal absorption is related to the rate of the compound crosses the intestinal wall to reach the portal blood circulation. The significant correlated relationships between NBE and blood-brain barrier penetration and between NBE and human intestinal absorption provide a simple but efficient metric to fastly judge the potential of a CSM into the brain from the circulation and that of a CSM into the circulation from the intestine.

In addition, we have similar and consistent observations on human metabolites. Interestingly, NBE distributions have bias in different cellular locations and in different body fluids, which could provide some valuable clues for metabolite-based drug discovery. In a summary, this study presented a simple but efficient metric to describe druggable properties of CSMs.NBE might provide greater helps when being combined with other in-silico methods or metrics in the future.

---

**Supplement Table 1. The list for the energy (KJ/mol) of various covalent bonds.**

## Notes

### Competing Interest Statement

The authors have declared no competing interest.

