## Supplement Table 1 for "A simple and efficient metric quantifying druggable property of chemical small molecules"

**Supplement Table 1**. The list for the energy (KJ/mol) of various covalent bonds[1]

| **Bond Type** | **Bond** | **Energy**(KJ/mol) |
|---|---|---|
| **Single Bond** | H—H | 432 |
| | H—F | 565 |
| | H—Cl | 427 |
| | H—Br | 363 |
| | H—I | 295 |
| | C—H | 413 |
| | C—C | 347 |
| | C—N | 305 |
| | C—O | 358 |
| | C—F | 485 |
| | C—Cl | 339 |
| | C—Br | 276 |
| | C—I | 240 |
| | C—S | 259 |
| | N—H | 391 |
| | N—N | 160 |
| | N—F | 272 |
| | N—Cl | 200 |
| | N—Br | 243 |
| | N—O | 201 |
| | O—H | 467 |
| | O—O | 146 |
| | O—F | 190 |
| | O—Cl | 203 |
| | O—I | 234 |
| | F—F | 154 |
| | F—Cl | 253 |
| | F—Br | 237 |
| | Cl—Cl | 239 |
| | Cl—Br | 218 |
| | Br—Br | 193 |
| | I—I | 149 |
| | I—Cl | 208 |
| | I—Br | 175 |
| | S—H | 347 |
| | S—F | 327 |
| | S—Cl | 253 |
| | S—Br | 218 |
| | S—S | 266 |
| | Si—Si | 340 |

| | | |
|---|---|---|
| | Si—H | 393 |
| | Si—C | 360 |
| | Si—O | 452 |
| **Double Bond** | C = C | 614 |
| | C = N | 615 |
| | O = O | 495 |
| | C = O | 745 |
| **Triple Bond** | C ≡ C | 839 |
| | C ≡ O | 1072 |
| | N = O | 607 |
| | N = N | 418 |
| | N ≡ N | 941 |
| | C ≡ N | 891 |

1. Bond Energies. Availabe online: https://chem.libretexts.org/Bookshelves/Physical_and_Theoretical_Chemistry_Textbook_Maps/Supplemental_Modules_(Physical_and_Theoretical_Chemistry)/Chemical_Bonding/Fundamentals_of_Chemical_Bonding/Bond_Energies (accessed on
